# Strong selective sweeps on the X chromosome in the human-chimpanzee ancestor explain its low divergence

**DOI:** 10.1101/011601

**Authors:** Julien Y. Dutheil, Kasper Munch, Kiwoong Nam, Thomas Mailund, Mikkel H. Schierup

## Abstract

The human and chimpanzee X chromosomes are less divergent than expected based on autosomal divergence. This has led to a controversial hypothesis proposing a unique role of the X chromosome in human-chimpanzee speciation. We study incomplete lineage sorting patterns between humans, chimpanzees and gorillas to show that this low divergence is entirely due to megabase-sized regions comprising one-third of the X chromosome, where polymorphism in the human-chimpanzee ancestral species was severely reduced. Background selection can explain 10% of this reduction at most. Instead, we show that several strong selective sweeps in the ancestral species can explain this reduction of diversity in the ancestor. We also report evidence of population specific sweeps in extant humans that overlap the regions of low diversity in the ancestral species. These regions further correspond to chromosomal sections shown to be devoid of Neanderthal introgression into modern humans. This suggests that the same X-linked regions that undergo selective sweeps are among the first to form reproductive barriers between diverging species. We hypothesize that meiotic drive is the underlying mechanism causing these two observations.

**Authors' Summary:** Because the speciation events that leads to human, chimpanzee and gorilla were close in time, their genetic relationship of these species varies along the genome. While human and chimpanzee species are most closely related, 15% of the human genome is more closely related to the gorilla genome than the chimpanzee genome, a phenomenon called incomplete lineage sorting (ILS). The amount and distribution of ILS can be predicted using population genetics theory, and is affected by demography and selection in the ancestral populations. It was previously reported that the X chromosome, in contrast to autosomes, is deprived of ILS, and this givies rise to controversial theories about the speciation event that splits humans and chimpanzees. Using a full genome alignment of the X chromosome, we show that this deprivation of ILS affects only one third of the chromosome. These regions also show reduced diversity in the extant populations of human and great apes, and coincide with regions devoid of Neanderthal introgression. We propose that these regions are targets of selection and that they played a role in the formation of reproductive barriers.

## Introduction

X chromosome evolution differs from that of autosomes in a variety of ways. The X chromosome is fully exposed to selection in males and is directly linked to the Y chromosome in male meiosis. Several recent studies in primates [1,2] and rodents [3] have shown that it experiences more adaptive evolution on the protein coding sequence than do the autosomes. Other studies have shown that the X chromosome has stronger and wider depressions in diversity around protein coding genes, which suggests that some combination of purifying and positive selection is more efficient than on autosomes [4–6].

The X chromosome also plays a special role in speciation by contributing disproportionately to hybrid incompatibility (the large X-effect) and it shows a stronger hybrid depression in males than in females (Haldane’s rule). A recent detailed investigation of introgression of Neanderthal genes into humans found that regions devoid of Neanderthal introgression are larger and more numerous on the X chromosome, suggestive of a role in reproductive isolation [7]. It has not been possible, however, to directly link these observations and the unique inheritance pattern of the X chromosome to speciation processes.

We and others have previously reported that (i) the average divergence of the human and chimpanzee X chromosomes is much lower than what would be expected from the autosomal divergence and (ii) that the X chromosome shows substantially less incomplete lineage sorting (ILS) between human, chimpanzee and gorilla than would be expected from the effective population size of the autosomes [8–10]. One hypothesis initially put forward by Patterson et al. [8] was that the speciation event of human and chimpanzee involved a secondary hybridization event after their initial split where most of the X chromosome of one species spread to both of the hybridizing species. Several authors have questioned this hypothesis [11–14] and it remains highly controversial.

Here we study the amount of incomplete lineage sorting between human, chimpanzee and gorilla along the X chromosome. We observe a striking pattern of mega-base sized regions with extremely low amounts of ILS, interspersed with regions with the amount of ILS expected from the effective population size of the X chromosome (that is, 3/4 that of the autosomes). We show that the most plausible explanation is several strong selective sweeps in the ancestral species to humans and chimpanzees. The low-ILS regions overlap strongly with regions devoid of Neanderthal ancestry in the human genome, which suggests that selection in these regions may create reproductive barriers. We suggest that the underlying mechanism is meiotic drive resulting from genetic conflict between the sex chromosomes, and that this is caused by testes expressed ampliconic genes found only on sex chromosomes and enriched in the regions where we find signatures of selective sweeps.

## Results

### Distribution of incomplete lineage sorting along the X chromosome

To explore the pattern of human-chimpanzee divergence across the full X chromosome we performed a detailed analysis of the aligned genomes of human, chimpanzee, gorilla and orangutan [10]. Using the coalescent hidden Markov model (CoalHMM) approach [15], we estimated demographic parameters in non-overlapping 1 Mb windows. For each window, we inferred the proportion of ILS using posterior decoding. The distribution of ILS proportions on autosomes follows a negatively skewed normal distribution (Figure 1A). The expected proportion of ILS in a 3-species alignment is given by the formula:

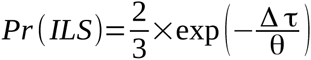

where Δ τ is the difference in speciation times and θ is the ancestral effective population size [16,17]. Estimates of these parameters from the gorilla genome consortium are Δ τ = 0.002468 and θ = 0.003232 [10]. From these parameters, the expected mean proportion of ILS should be 31.06%, in agreement with the observed 30.58%.

**Fig. 1.**
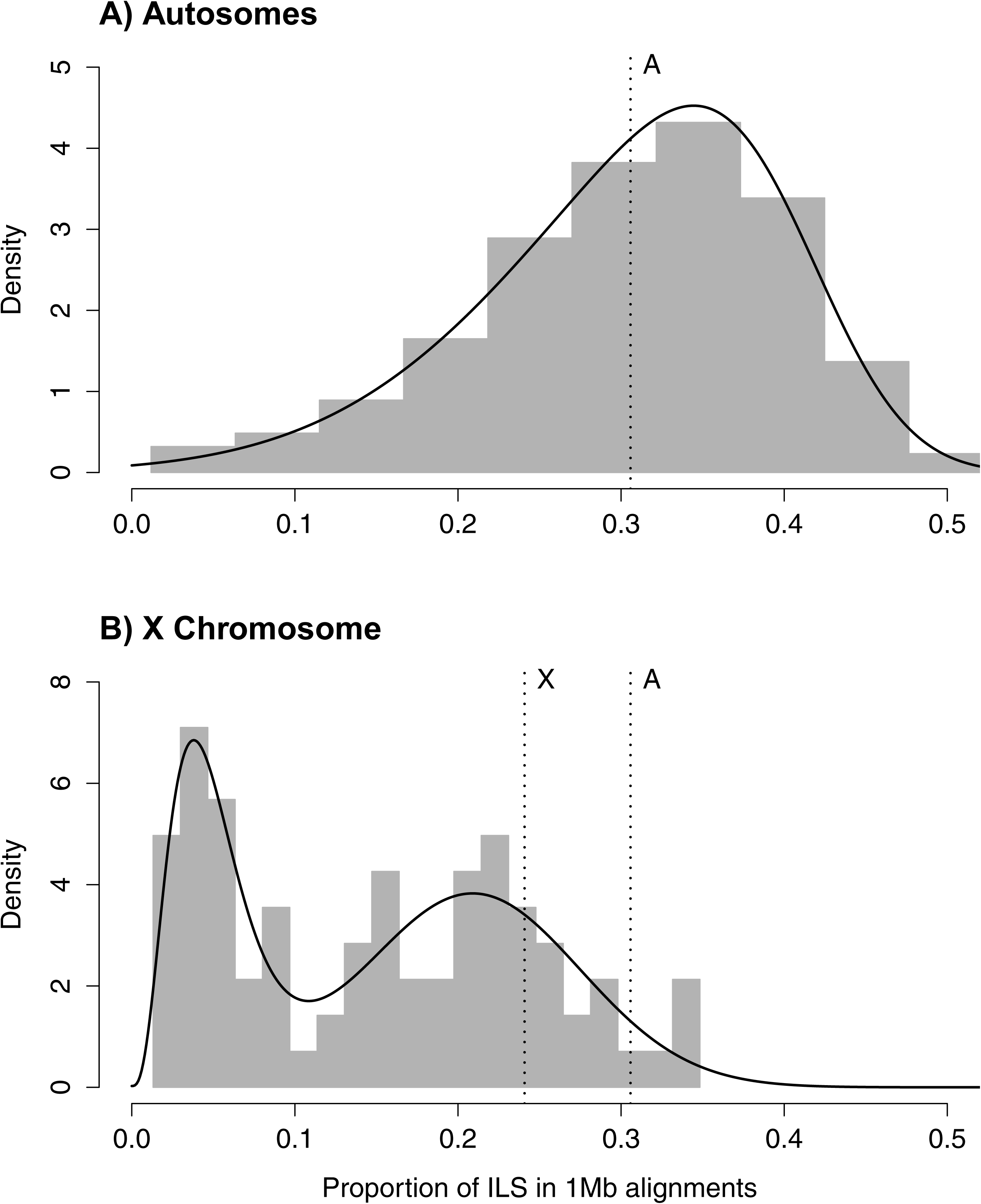
Distribution of incomplete lineage sorting (ILS) along the Human genome for autosomes (A) and the X chromosome (B). Grey bars show the distribution of ILS as estimated from the posterior decoding of the CoalHMM model. Solid black lines show the best fit of a skewed normal distribution in (A) and a mixture of a Gamma and a Gaussian distribution in (B). The A-labeled vertical line show the median of ILS on the autosomes (A), reported on the X chromosome (B). The X-labeled vertical line shows the expectation of ILS on the X chromosome based on the estimate of ILS on the autosomes. The second mode of the distribution of ILS on the X chromosome matches this expectation.

Assuming that the ancestral effective population size of the X chromosome, θ_*X*_, is three quarters that of the ancestral effective population size of the autosomes, the expected amount of ILS on the X chromosome should be 24.08%. The distribution of ILS proportions on the X chromosome is bimodal (Figure 1B) and in stark contrast to the distribution on the autosomes. One mode represents 63% of the alignment, with a mean proportion of ILS of 21%, close to the expectation of 24% (the 99% confidence interval of the high ILS mode is [17.6%, 24.5%], estimated using parametric bootstrap). The second mode is estimated to represent 37% of the alignment and shows a mean proportion of ILS below 5%. The regions exhibiting low ILS form 8 major segments spread across the X chromosome (Table 1 and Figure 2A) and cover 29 Mb out of a total alignment length of 84 Mb. Region X5 is split in two by the centromeric region, where alignment data are missing. These striking patterns suggest that unique evolutionary forces have shaped the ancestral diversity in these regions.

**Table 1.** Low-ILS regions on the X chromosome. Coordinates are given according to the Human genome hg19.

| Region | Begin | End | Average ILS |
| --- | --- | --- | --- |
| X1 | 10,241,177 | 12,619,185 | 0.035 |
| X2 | 16,946,047 | 18,747,389 | 0.054 |
| X3 | 19,303,480 | 22,198,160 | 0.047 |
| X4 | 38,344,992 | 41,272,675 | 0.062 |
| X5 | 45,930,478 | 77,954,462 | 0.050 |
| X6 | 99,459,295 | 111,145,964 | 0.031 |
| X7 | 128,232,540 | 136,796,526 | 0.034 |
| X8 | 151,519,514 | 155,156,362 | 0.050 |

**Fig. 2.**
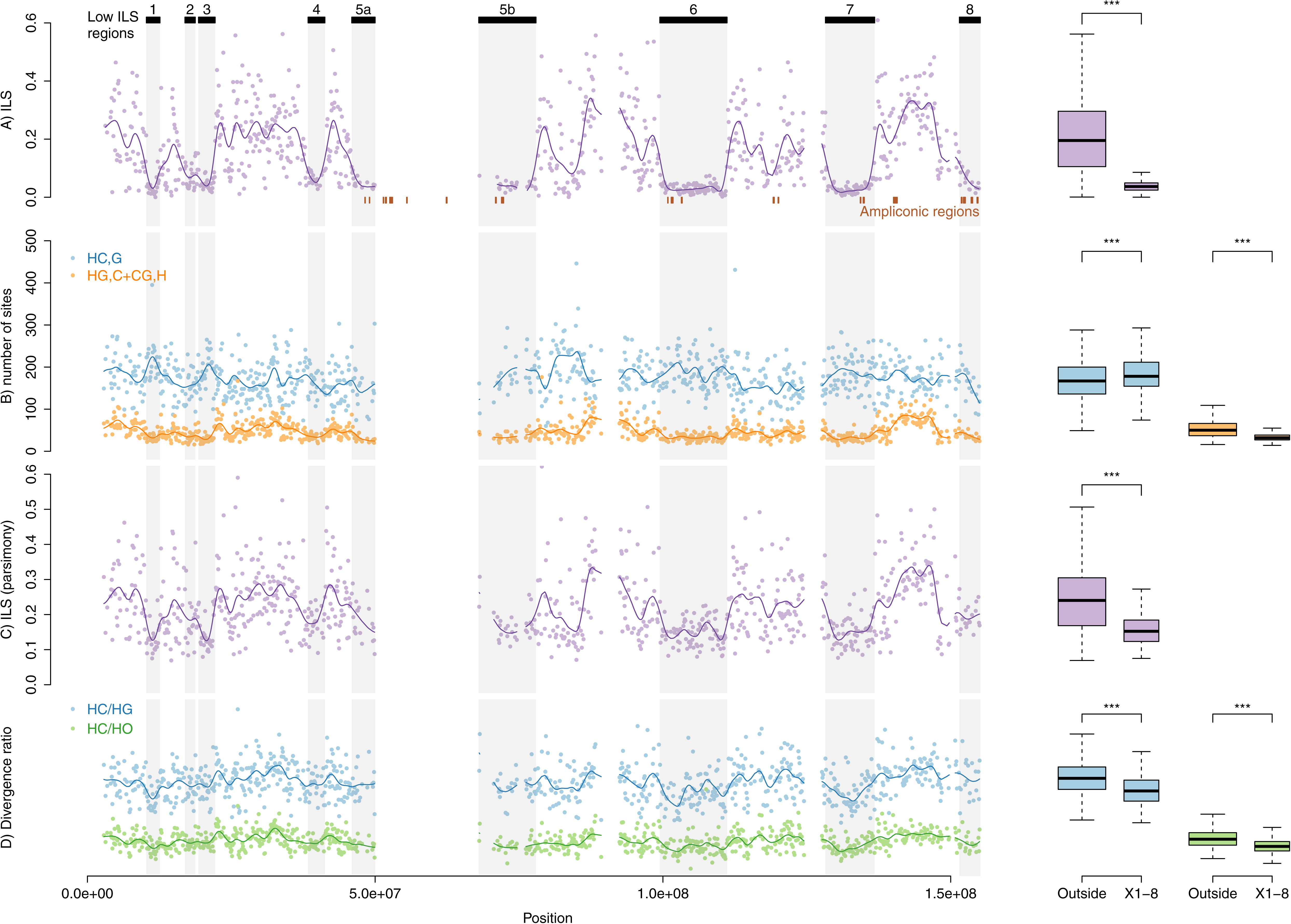
Patterns of incomplete lineage sorting along the X chromosome. Graphs on the left show variation along the chromosome, graphs on the right contrast the distribution of low-ILS regions version the rest of the chromosome. Significance codes are according to Wilcoxon rank test. Rows: (A) Proportion of inferred ILS in individual non-overlapping 100 kb windows and a fitted spline. Inferred regions with low ILS are shown on top, and reported on all figures. (B) Frequencies of parsimony informative sites in 100 kb windows, supporting both the canonical genealogy (HC),G and the alternative ones (HG),C and (CG),H together. (C) ILS as estimated by the proportion of parsimony informative sites supporting an alternative topology. D) Ratio of divergences HC/HG and HC/HO estimated in 100 kb windows.

### Robustness of ILS estimation

In Scally et al. [10], we independently estimated parameters in non-overlaping windows of 1 Mb, allowing for parameters to vary across the genome. To test whether inference of very low proportions of ILS could result from incorrect parameter estimation, we compared the inferred amount of ILS under alternative parameterizations with that inferred using fixed parameters (either all or speciation time parameters only) along the genome. These alternative parametrizations result in very similar estimates of ILS (Figure 3 and corresponding UCSC genome browser tracks at http://bioweb.me/HCGILSsupp/UCSCTracks/).

**Fig. 3.**
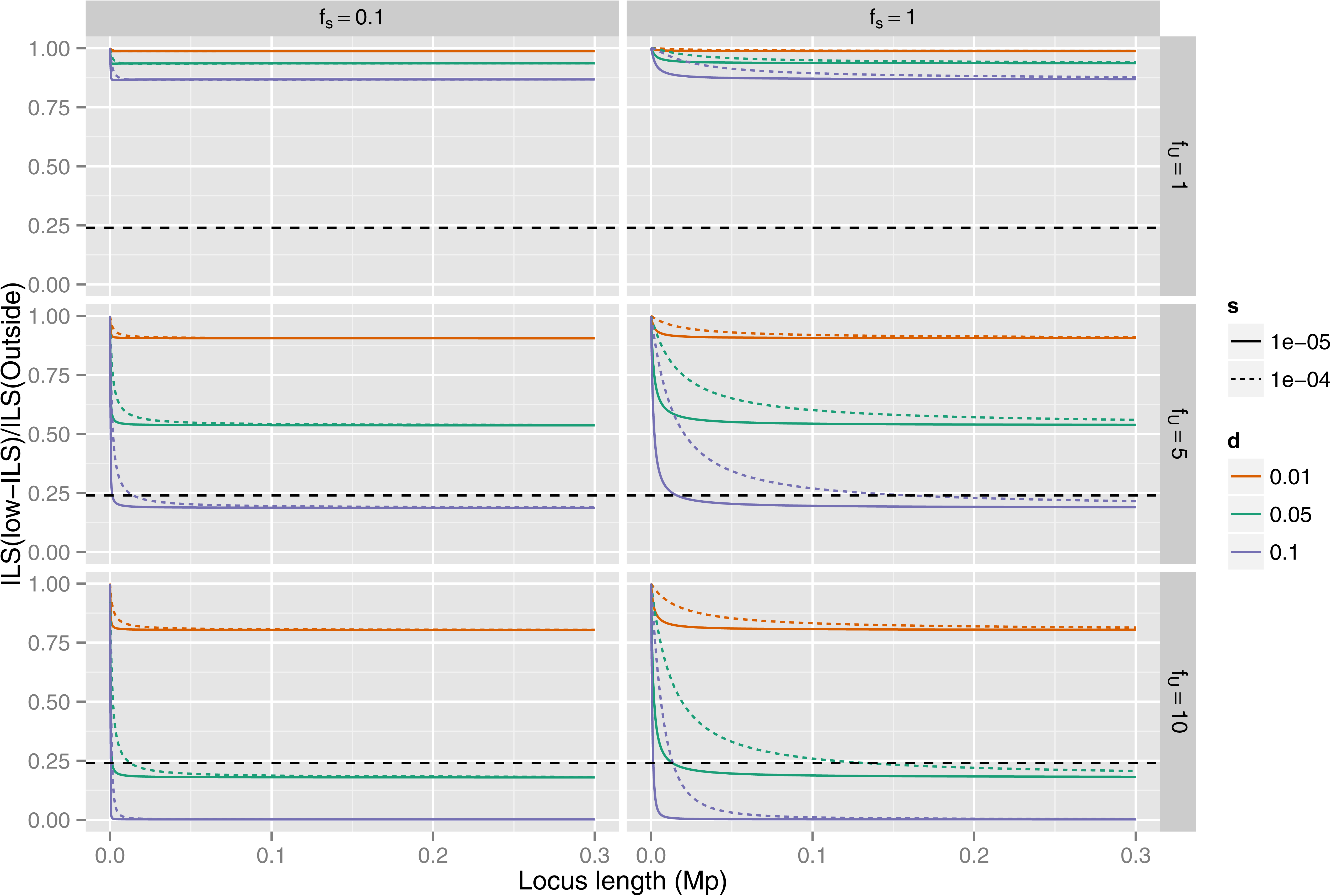
Background selection and ILS. The plots show the ratio of ILS inside the low ILS regions compared to that outside the regions, assuming speciation times of 5.95 mya and 3.7 mya, 20 year generations and that the neutral X effective population size is 3/4 that of the autosomes. The colors corresponds to different choices of which fraction of mutations are deleterious, varying from 1% to 10%. The different columns correspond to different choices of selection within the low ILS regions – set to either the same as outside or one tenth of the selection strength outside – and different rows show how much more of the regions is under selection compared to outside, either the same or a factor of five or ten. Selection strength is set to either 1e-4 (dotted curve) or 1e-5 (solid curve). The horizontal dashed line represents the observed reduction in ILS of 24% (from 21% ILS outside low-ILS regions to the <5% ILS of low-ILS regions).

Our observations do not reflect a lower power to detect ILS in the identified regions. To address this possibility, we counted the number of informative sites supporting each of the three alternative topologies connecting humans, chimpanzees and gorillas in non-overlapping 100 kb windows along the alignment. While the total frequency of parsimony-informative sites is significantly lower in the low-ILS regions compared with the rest of the genome (0.00270 vs. 0.00276, Fisher’s exact test p-value = 1.34e-05), there is a highly significant excess of sites supporting the species topology (0.00229 vs. 0.00210, Fisher’s exact test p-value < 2.2e-16) and deficit of sites in these regions supporting ILS topologies (0.00042 vs. 0.00066, Fisher's exact test p-value < 2.2e-16, Figure 2B-C), consistent with a lower proportion of ILS.

We computed the ratio of human-chimpanzee divergence to human-gorilla divergence and human-orangutan divergence in 100 kb windows. Assuming a constant mutation rate across the phylogeny and constant ancestral effective population sizes along the genome, these ratios should remain constant. However, the low-ILS regions show a relatively lower human-chimpanzee divergence. This is expected based on a lower ancestral diversity of the human-chimpanzee ancestor in these regions (Figure 2D). A lower mutation rate in these regions would explain this pattern only if the reduction is restrained to the human-chimpanzee lineage.

### The effect of background selection on ILS

Deleterious mutations are continuously pruned from the population through purifying selection, reducing the diversity of linked sequences. Such background selection potentially plays an important role in shaping genetic diversity across the genome [18]. The strength of background selection increases with the mutation rate, with density of functional sites, with decreasing selection coefficient against deleterious mutations and with increasing recombination rate [19]. Low-ILS regions display both a 0.6-fold lower recombination rate compared to the rest of the chromosome (1.01 cM/Mb versus 1.62 cM/Mb, Wilcoxon test p-value = 2.2e-07) as well as a two-fold higher gene density - a proxy for the proportion of functional sites (3.1% exonic sites versus 1.5% on average, Wilcoxon test p-value < 2.2e-16). Background selection is therefore both expected to be more common (by a factor of ∼2.1 due to more functional sites) and to affect larger regions (by a factor of ∼1.8 due to less recombination) in the low-ILS regions. To estimate extent to which this may explain our observations, we used standard analytical results that estimate the combined effect of multiple sites under purifying selection (see Material and Methods). Even if we assume that the proportions of functional sites in the candidate regions is two times higher than the observed number of exon base pairs, and that all mutations at these sites are deleterious with a selection coefficient that maximizes the effect of background selection, the expected proportion of ILS should only be reduced by approximately 10% relative to the level found on the remaining X chromosome (19% ILS compared to 21% ILS). To explain the observed reductions in ILS by background selection alone, unrealistic differences of functional site densities are required (*e.g*. 50% inside identified regions and 10% outside, see Figures 3 and S2). As a further line of evidence, we computed the maximal expected reduction of ILS based on the observed density of exonic sites and average recombination rate (see Methods). We find that only 79 of 252 analyzable windows (31%) could be explained by the action of background selection only, an observation incompatible with the hypothesis that background selection is the sole responsible for the widespread reduction of ILS along the X chromosome.

Finally, background selection is expected to have a stronger effect on the autosomes than on the X chromosome. The fact that we do not observe large regions devoid of ILS on the autosomes further argues against background selection as the major force creating the observed large regions with reduced ILS on the X chromosome.

### Selective sweeps and ILS

Adaptive evolution may also remove linked variation during the process of fixing beneficial variants. In the human-chimpanzee ancestor, such selective sweeps will have abolished ILS at the locus under selection and reduced the proportion of ILS in a larger flanking region. Several sweeps in the same region can this way result in a strong reduction of ILS on a mega-base scale. We simulated selective sweeps in the human-chimpanzee ancestor using a rejection sampling method (see Material and Methods). A single sweep is only expected to reduce ILS to less than 5% on a mega-base wide region if selection coefficients are unrealistically high (s > 0.2), suggesting that several sweeps have contributed to the large-scale depletions of ILS (Figures 4 and S3).

**Fig. 4.**
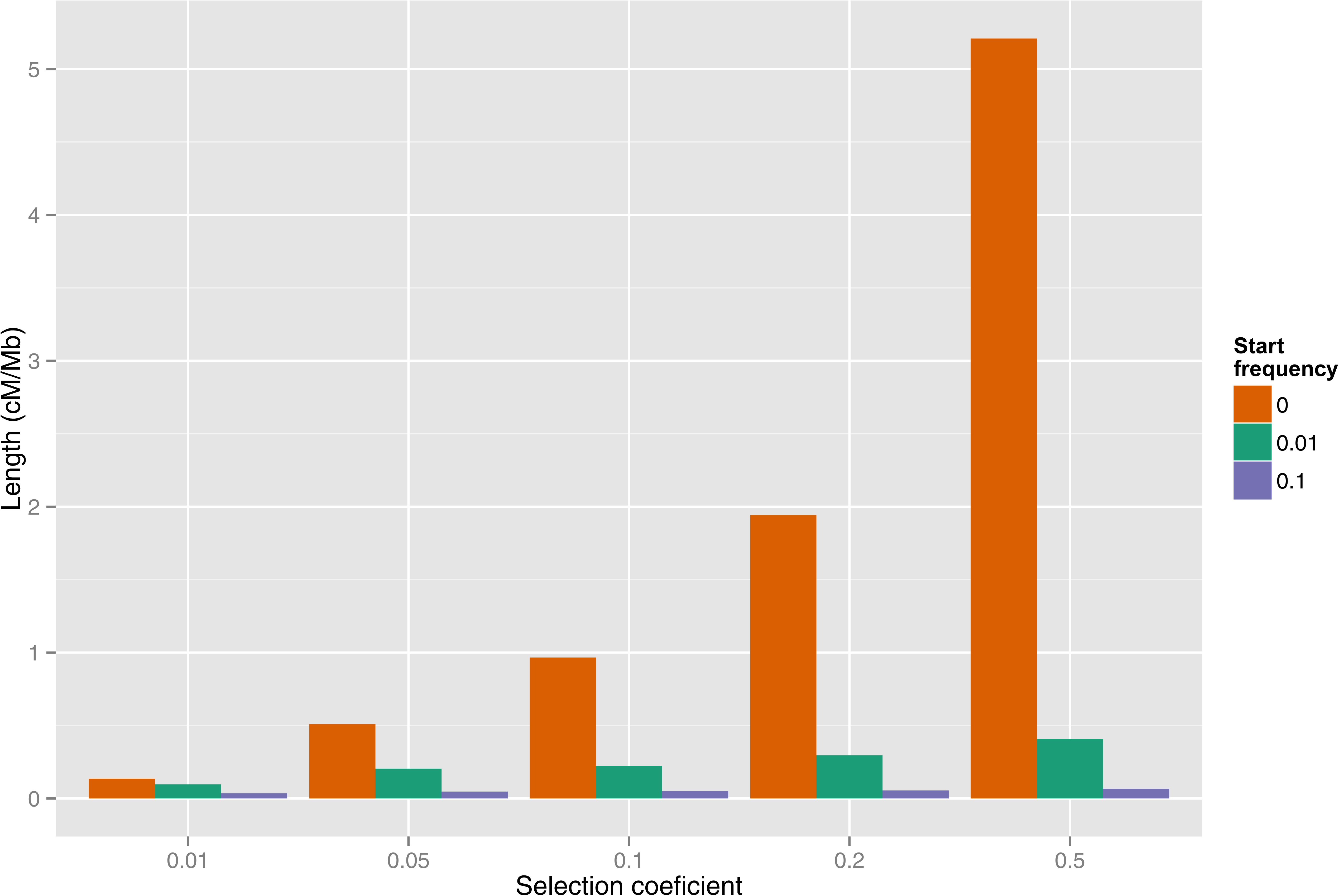
Expected genetic length of the region with less than 5% ILS surrounding a selected mutant with given selection coefficient and start frequency.

If the low-ILS regions are indeed subject to recurrent sweeps, they are expected to also show reduced diversity in human populations. We therefore investigated the patterns of nucleotide diversity in the data of the 1000 Genomes Project [20]. We computed the nucleotide diversity in 100 kb non-overlapping windows along the X chromosome and compared windows within and outside low-ILS regions. Figure 5 summarizes the results for the CEU, JPT and YRI populations (results for all populations are shown in Figure S4). We find that diversity is significantly reduced in all low-ILS regions compared with the chromosome average (Table 2), and this reduction is on average significantly greater in the Asian and European populations than in the African population (analysis of variance, see Material and Methods). The same analysis was performed on each of the eight low-ILS regions independently, and revealed differences between regions (Table 3). Plotting population specific diversity across the X chromosome reveals several cases of large-scale depletions of diversity in both Europeans and East Asians. While these depletions affect similar regions, their width differ between populations. This finding suggests that strong sweeps in these regions occurred independently in the European and East Asian population after their divergence less than 100,000 years ago.

**Table 2.** Reduction of diversity (measured in 100 kb non-overlapping windows) in low-ILS regions in Human populations as compared to the X chromosome mean outside the low-ILS regions. Stars denote significance of p-values of Wilcoxon tests corrected for multiple testing: 10% (.), 5% (*), 1%(**) < 1% (***).

| Population | Region 1 | Region 2 | Region 3 | Region 4 | Region 5 | Region 6 | Region 7 | Region 8 |
| --- | --- | --- | --- | --- | --- | --- | --- | --- |
| GBR | 73% (*) | 36% (***) | 42% (***) | 79% (*) | 48% (***) | 50% (***) | 53% (***) | 65% (**) |
| FIN | 78% (.) | 35% (***) | 45% (***) | 81% (.) | 48% (***) | 47% (***) | 54% (***) | 59% (***) |
| CHS | 72% (*) | 49% (***) | 32% (***) | 77% (.) | 47% (***) | 50% (***) | 67% (***) | 72% (*) |
| PUR | 78% (*) | 40% (***) | 56% (***) | 81% (*) | 58% (***) | 51% (***) | 54% (***) | 68% (***) |
| CLM | 75% (*) | 43% (***) | 47% (***) | 76% (*) | 55% (***) | 54% (***) | 58% (***) | 70% (**) |
| IBS | 71% (*) | 41% (***) | 39% (***) | 84% (NS) | 48% (***) | 53% (***) | 52% (***) | 55% (***) |
| CEU | 73% (*) | 36% (***) | 39% (***) | 78% (*) | 51% (***) | 47% (***) | 54% (***) | 62% (***) |
| YRI | 79% (*) | 52% (***) | 64% (***) | 78% (**) | 60% (***) | 66% (***) | 56% (***) | 70% (***) |
| CHB | 73% (*) | 45% (***) | 29% (***) | 75% (*) | 46% (***) | 50% (***) | 66% (***) | 70% (*) |
| JPT | 76% (.) | 47% (***) | 32% (***) | 81% (NS) | 46% (***) | 46% (***) | 66% (***) | 67% (*) |
| LWK | 79% (*) | 52% (***) | 65% (***) | 80% (**) | 63% (***) | 65% (***) | 57% (***) | 67% (***) |
| ASW | 77% (*) | 50% (***) | 65% (***) | 77% (**) | 65% (***) | 65% (***) | 54% (***) | 69% (***) |
| MXL | 79% (.) | 43% (***) | 39% (***) | 83% (.) | 58% (***) | 53% (***) | 54% (***) | 68% (**) |
| TSI | 80% (.) | 35% (***) | 42% (***) | 76% (*) | 50% (***) | 51% (***) | 55% (***) | 60% (***) |

**Fig. 5.**
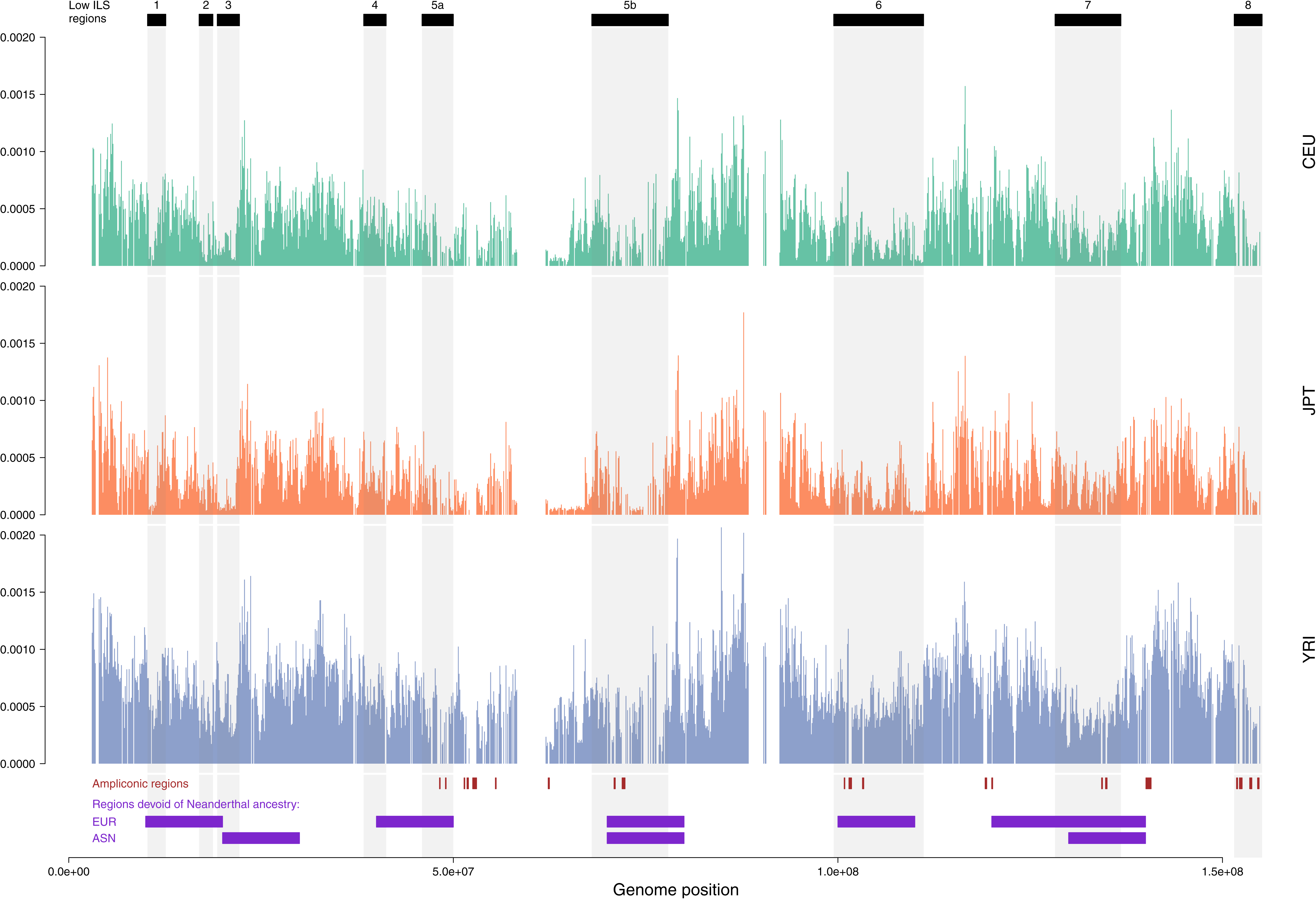
Distribution of nucleotide diversity along the X chromosome of human populations. Nucleotide diversity is computed in 100 kb non-overlapping windows. Ampliconic regions [24] as well as regions absent of Neanderthal introgression [7] are shown at the bottom. Figure S4 shows all 14 populations.

**Table 3.** Average reduction of diversity for each population group and low-ILS region. For each region, populations with the same letter code are not significantly different according to Tukey's posthoc test (5% level).

| Population | Total | Region 1 | Region 2 | Region 3 | Region 4 | Region 5 | Region 6 | Region 7 | Region 8 |
| --- | --- | --- | --- | --- | --- | --- | --- | --- | --- |
| Africa | 64% (a) | 78% (a) | 51% (a) | 64% (a) | 78% (a) | 63% (a) | 65% (a) | 55% (a) | 70% (a) |
| America | 57% (b) | 77% (a) | 42% (ab) | 47% (b) | 80% (a) | 57% (b) | 53% (b) | 55% (a) | 70% (a) |
| Asia | 53% (c) | 74% (a) | 47% (a) | 31% (c) | 78% (a) | 46% (c) | 49% (c) | 67% (b) | 70% (ab) |
| Europe | 53% (c) | 75% (a) | 37% (b) | 41% (b) | 80% (a) | 49% (d) | 50% (c) | 54% (a) | 60% (b) |

## Discussion

Using a complete genome alignment of human, chimpanzee, gorilla and orangutan, we report that the human-chimpanzee divergence along the X chromosome is a mosaic of two types of region: two thirds of the X chromosome display a divergence compatible with the expectation of an ancestral effective population size of the X equal to three quarters that of the autosome, while one third of the X chromosome shows an extremely reduced divergence, and is virtually devoid of incomplete lineage sorting. We have demonstrated that such desert of diversity cannot be accounted for by background selection alone, but must result from recurrent selective sweeps.

If the low-ILS regions evolve rapidly through selective sweeps, they could be among the first to accumulate hybrid incompatibility between diverging populations. Recently, the X chromosome was reported to exhibit many more regions devoid of Neanderthal introgression into modern humans than the autosomes. This suggests an association of negative selection driven by hybrid incompatibility with these X-linked regions [7]. We find a striking correspondence between regions of low ILS and the regions devoid of Neanderthal introgression (Figures 2 and 5). We recently reported dramatic reductions in X chromosome diversity in other great ape species that almost exclusively affect areas of the low-ILS regions [21] (see Figure S5). Taken together, these findings show that the regions on the X chromosome that contributed to hybrid incompatibility in the secondary contact between humans and Neanderthals have been affected by recurrent, strong selective sweeps in humans and other great apes. The occurrence of a secondary contact between initially diverged human and chimpanzee populations (the complex speciation scenario of Patterson *et al* [8]) is therefore compatible with a lower proportion of ILS in these regions. In this scenario the depletions of ILS would result from negative selection leading to the fixation of large genomic regions contributed from by only one of the admixing populations, as suggested by Sankararaman *et al [7]*.

However, such complex speciation scenarios do not explain the large-scale reductions of diversity in extant species. We propose a hypothesis that may better account for the generality of our findings: Deserts of diversity may arise via meiotic drive, through which fixation of variants that cause preferential transmission of either the X or Y chromosome produces temporary sex ratio distortions [22]. When such distortions are established, mutations conferring a more even sex ratio will be under positive selection. Potential candidates involved in such meiotic drive are ampliconic regions, which contain multiple copies of genes that are specifically expressed in the testis. These genes are postmeiotically expressed in mice, and a recent report suggests that the Y chromosome harbors similar regions [23]. Fourteen of the regions identified in humans [24] are included in our alignment, 11 of which are located in low-ILS regions (Figure 2), representing a significant enrichment (binomial test with p-value = 0.0019). Whatever the underlying mechanism, our observations demonstrate that the evolution of X chromosomes and their role in speciation merits further study.

## Material and Methods

### Genome alignment and data pre-processing

The Enredo/Pecan/Ortheus genome alignment of the five species human, chimpanzee, gorilla, orangutan and macaque from [10] was used as input. In order to remove badly sequenced and / or ambiguously alignment regions, we filtered the input 5-species alignments using the MafFilter program [25]. We sequentially applied several filters to remove regions with low sequence quality score and high density of gaps. Details on the filters used can be found in the supplementary material of Scally et al. [10].

### Inference of incomplete lineage sorting

The divergence of two genomes depends on both the mutation rate and underlying demographic scenario. With a constant mutation rate μ and simple demography (constant sized panmictic population evolving neutrally), the time to the most recent common ancestor of two sequences sampled from different species is given by a constant species divergence, τ=*T*. μ T, and an ancestral coalescence time following an exponential distribution with mean θ=2. *Ne*_*A*_. μ, where T is the number of generations since species divergence and Ne_A_ is the ancestral effective population size [9,26]. For species undergoing recombination, a single individual genome is a mosaic of segments with distinct histories, and therefore displays a range of divergence times [8,9,27]. When two speciation events separating three species follow shortly after each other, this variation of genealogy can lead to incomplete lineage sorting (ILS), where the topology of gene trees do not correspond to that of the species tree [9,16]. Reconstructing the distribution of divergence along the genome and the patterns of ILS allows inference of speciation times and ancestral population sizes. We used the CoalHMM framework to infer patterns of ILS along the X chromosome. Model fitting was performed as described in [10]. ILS was estimated using posterior decoding of the hidden Markov model as the proportions of sites in the alignment which supported one of the (HG),C or (CG),H topologies.

### Distribution of ILS

For the autosomal distribution of ILS, we fitted a skewed normal distribution (R package ‘sn' [28]) using the *fitdistr* function from the MASS package for R. For the X chromosome ILS distribution, we fitted a mixture of gamma and Gaussian distribution. The mixed distribution follows a normal density with probability p, and a gamma density with probability 1-p. In addition to p, the mixed distribution has four parameters: the mean and standard deviation of the Gaussian component, and the shape and rate of the gamma component. The L-BFGS-B optimization method was used to account for parameter constraints. Resulting parameter estimates are 0.209 for the mean of the Gaussian component, 0.066 for the standard deviation of the Gaussian component, 4.139 for the alpha parameter (shape) of the gamma component, 83.369 for the beta parameter (rate) of the gamma component, and p = 0.632. The mean of the gamma component is alpha / beta = 0.0497, that is, less than 5% ILS. We compared the resulting fit with a mixture of skewed normal distributions, which has two extra parameters compared to a Gamma-Gaussian mixture, and found that the skew of the higher mode is very close to zero, while the Gamma distribution offered a better fit of the lower mode. We used a parametric bootstrap approach to estimate the confidence interval of the proportion of ILS for the mean of the normal component of the mixed distribution. We generated a thousand pseudo-replicates by sampling from the estimated distribution, and we re-estimated all parameters from each replicate in order to obtain their distribution. Replicates where optimization failed were discarded (40 out of 1000).

### Characterization of low-ILS regions

In order to characterize the patterns of ILS at a finer scale, we computed ILS in 100 kb windows sliding by 20 kb along the alignment. To exhibit regions devoid of ILS, we selected contiguous windows with no more than 10% of ILS each. Eight of these regions were greater than 1 Mb in size, and their resulting amount of ILS is less than 5% on average (Table 1). The coordinates of these regions were then translated according to the human hg19 genome sequence. These data are available as a GFF file for visualization in the UCSC genome browser at http://bioweb.me/HCGILSsupp/.

### Reduction in ILS by background selection

Background selection reduces diversity by a process in which deleterious mutations are continuously pruned from the population. The strength of background selection in a genomic region is determined by the rate at which deleterious mutations occur, *U*, the recombination rate of the locus, *R*, and the strength of negative selection on mutants, *s*. We consider the diversity measure, π (the pairwise differences between genes) which in a randomly mating population is linearly related to the effective population size. If π_0_ denotes diversity in the absence of selection and π the diversity in a region subject to background selection, then the expected reduction in diversity is given by

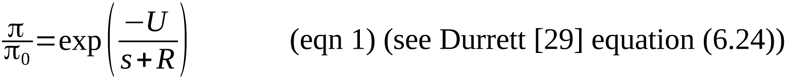

The rates *U* and *R* are both functions of the locus length (*U*=*uL* and *R*=*rL*) where *r* denotes the per-nucleotide-pair recombination rate, *u* the per-nucleotide deleterious rate, and *L* the length of the locus. To investigate if background selection can explain the observed reductions in ILS we must compute the expected reduction in diversity in the low-ILS regions relative to the reduction in the remaining chromosome. A larger reduction in low-ILS regions may be caused by weaker negative selection, higher mutation rate, lower recombination rate, and larger proportion of functional sites at which mutation is deleterious. To model the variation of these parameters inside and outside lowILS regions we simply add a factor to each relevant variable. The relative reduction can thus be expressed as:

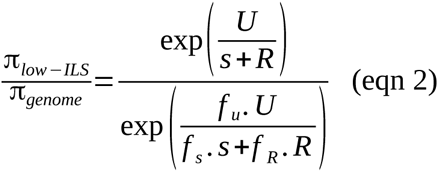

The recombination rate, *R*, and the factor, *f*_*R*_, can be obtained from the deCODE recombination map [30]. We computed the average deCODE recombination rate, as well as the proportion of sites in exons (as a measure of selective constraint) in non-overlapping 100 kb along the human X chromosome.

The recombination rate average outside the low ILS regions is 1.62 cM/Mb and the recombination rate inside the regions is 1.01 cM/Mb which gives us *f*_*R*_ =0.6. For the remaining parameters, *s* and *U*, we need to identify realistic values outside the lowILS regions. Background selection is stronger when selection is weak, but the equation is not valid for very small selection values where selection is nearly neutral. Once *s* approaches 1/ *N*_*e*_, we do not expect any background selection. Most estimates of effective population sizes, *N*_*e*_, in great apes are on the order 10,000-100,000 and this puts a lower limit on relevant values of *s* at 10^-4^ - 10^-5^. To conservatively estimate the largest possible effect of background selection we explore this range of selection coefficients: *s*=10^−4^ and *s*=10^−5^ and allow the selection inside the low ILS regions to be one tenth (*f*_*S*_=0.1) of that outside. For *U* values outside low-ILS regions we assume the mean human mutation rate, estimated to be 1.2⋅10^−8^ per generation [31]. To obtain the rate of deleterious mutation we must multiply this with the proportion of sites subject to weak negative selection, *d*. Although this proportion is subject to much controversy it is generally believed to be between 3% and 10% [32]. However, as explained below we explore values up to 100% inside the low-ILS regions.

We assessed the relative diversity for combinations of *s* and *d* values (Figure S2). Each cell represents a combination of parameter values for *s*, *d*, *f*_*U*_ and *f*_*s*_. The reduction of diversity Δπ translates into reduction of ILS, Δ *ILS* (Figure 3). Assuming the time between speciation events, the generation time and population size reported in Scally et al. [10] (Δ *T* = 2,250,000 years, g = 20) ILS is given by

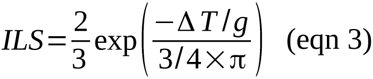

and the relative ILS is given by

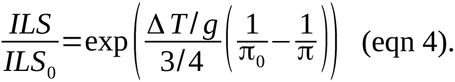

For the most extreme parameter values, we see a relative reduction in ILS of nearly 100%. In these cases, however, 100% of the nucleotides within low-ILS regions are under selection. In the cases where 25% of the nucleotides in the low-ILS regions are under selection compared to 5% outside (*f*_*U*_ =5, *d*=0.05), the regions retain more than half of the diversity seen outside the regions.

We further computed the expected reduction of ILS due to background selection in 100 kb windows located in low-ILS regions using equation (4). For each window, we computed the frequency of sites in exons and the average DECODE recombination rate. We further assumed a selection coefficient *s*=10^−5^ and allow the selection inside the low ILS regions to be one tenth (*f*_*S*_=0.1). Out of 285 windows located in low-ILS regions, we could estimate the maximal reduction of ILS due to background selection in 252 windows for which a DECODE recombination estimate was available. In 79 of these windows the expected reduction was below 0.20.

### Simulation of ancient selective sweeps

To assess how hard and soft sweeps in the human-chimpanzee ancestor can have reduced the proportion of ILS we simulated sweeps for different combinations of selection coefficients, s, and frequencies of the selected variant at the onset of selection, *f*. Frequency trajectories of selected variants are obtained using rejection sampling to obtain trajectories that fix in the population. Trajectories used to simulate hard sweeps begin at one and proceed to fixation at 2N * 3/4 by repeated binomial sampling with probability parameter N_mut_/(N_mut_ + (N - N_mut_)(1-s)), where N_mut_ is the number of selected variants in the previous generation. We use a human-chimpanzee speciation time of 3.7 Myr, a human-gorilla speciation time of 5.95 Myr, a human-chimpanzee effective population size of 73,200 as reported in [10], assuming a mutation rate of 1e-9 and a generation time of 20 years. Trajectories used to simulate soft sweeps are constructed by joining two trajectories. If f is the frequency of the variant at the onset of selection F= f * 2N * 3/4 is the number of variants. We first sample a trajectory that represents the time before the onset of selection. This trajectory is required to reach F at least once before it fixes or is lost, and is truncated randomly at one of the points where it passes the value F. The truncated trajectory is then appended with a trajectory under selection that begins at F and proceeds to fixation.

In each simulation we consider a sample of two sequences that represent 10 cM. As the effect of the sweep is symmetric we only simulate one side of the sweep. We then simulate backwards in the Wright-Fisher process with recombination allowing at most one recombination event per generation per lineage but allowing mergers of multiple lineages expected to occur in strong sweeps. The simulation proceeds until all sequence segments have found a most recent common ancestor (TMRCA). For each combination of parameters s and f we perform 1,000 simulations and the mean TMRCA is computed in bins of 10 kb.

In each simulation individual sequence segments are called as ILS with probability 2/3 if the TMRCA exceeds the time between the speciation events. The width of the region showing less than 5% ILS is then computed for each simulation. In Figure 4 and S3 a recombination rate of 1 cM/Mb is assumed to translate to physical length.

### Comparing diversity between human populations

We computed the nucleotide diversity in 100 kb non-overlapping windows along the X chromosome for the 14 populations from the 1,000 genomes project. The windows in each low-ILS region were compared to windows outside the regions using a Wilcoxon test with correction for multiple testing [33] (Table 2). We computed the relative nucleotide diversity in the 1,298 windows located in low-ILS regions by dividing by the average of the rest of the X chromosome. Each population was further categorized according to its origin, Africa, America, Asia or Europe [20]. A linear model was fitted after Box-Cox transformation:

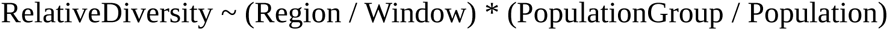

where Window is the position of the window on the X chromosome, and is therefore nested in the (low-ILS) Region factor. Analysis of variance reveals a highly significant effect of the factors Region and Window (p-values < 2e-16), PopulationGroup (p-value < 2e-16) and their interactions (p-value < 2e-16). The nested factor Population however was not significant, showing that the patterns of relative diversity within low-ILS regions are similar between populations within groups. A Tukey's Honest Significance Difference test (as implemented in the R package 'agricolae') was performed on the fitted model and further revealed that European and Asian diversity are not significantly different, while they are from African and American diversity.

### Association with ampliconic regions

The coordinates of ampliconic regions tested in [24] were translated to hg19 using the liftOver utility from UCSC. Fourteen regions were included in our alignment. Eleven regions have a midpoint coordinate within a low-ILS region. With 37% of the positions on the X being within a low-ILS region, a unilateral binomial test leads to a p-value = 0.001879501, meaning that the observed proportion of ampliconic regions within low-ILS regions is significantly higher than expected by chance.

## Acknowledgments

The authors would like to thank David Reich, Nick Patterson and Thomas Bataillon for discussions, and Sriram Sankararaman for sharing the coordinates of the regions devoid of Neanderthal ancestry. JYD acknowledges the LOEWE-Zentrum für Synthetische Mikrobiologie (Synmikro) for funding. KM and TM are funded by the Danish Council for Independent Research. This publication is the contribution no. XXXX-XXX of the Institut des Sciences de l'Évolution de Montpellier (ISE-M).

## Author Contributions

JYD, KM, TM and MHS designed the study and wrote the manuscript. JYD performed the incomplete lineage sorting analysis. JYD and KN analyzed data from the 1000 Genome Project. KM and TM performed calculations and simulations for background selection and selective sweeps. KN analyzed data from great apes.

## Supplementary figures

**Fig. S1.** Effect of parameter estimation on ILS inference on the X chromosome alignment. ILS is computed in 1 Mb alignments. The x-axis shows the inferred amount of ILS when model parameters are estimated independently on each alignment (free parameters). The left graph shows the amount of ILS inferred when all model parameters are assumed constant along the X chromosome, estimated from the full chromosome alignment (fixed parameters). The right graph shows the amount of ILS inferred when only the speciation times are considered constant along the chromosome; ancestral population sizes and recombination rate are allowed to vary and are estimated independently for each alignment.

**Fig. S2.** Background selection and diversity. The plots show the ratio of nucleotide diversity inside the low ILS regions compared to that outside the regions, assuming speciation times of 5.95 mya and 3.7 mya, 20 year generations and that the neutral X effective population size is 3/4 that of the autosomes. Rest of legend is as in Figure 3.

**Fig. S3.** Distribution of the genetic length of the region with less than 5% ILS extending away from a selected mutant. Each panel shows the distribution for a combination of selection coefficient, and frequency of the mutant at the onset of selection. Each sub-plot is based on 1,000 simulations.

**Fig. S4.** Distribution of nucleotide diversity along the X chromosome for the 14 populations from the 1000 Genomes Project. Nucleotide diversity is computed in 100 kb non-overlapping windows. Ampliconic regions [24] as well as regions absent of Neanderthal introgression [7] are shown at the bottom.

**Fig. S5.** Nucleotide diversity of 100 kb windows in low-diversity regions (< 20% of species average) in great apes. Blue bars represent low-ILS regions identified in this study. B: Bonobo, CC: Central chimpanzee, EC: Eastern chimpanzee, WC: Western chimpanzee, NC: Nigerian chimpanzee, WLG: Western lowland gorilla, SO: Sumatran orangutan, BO: Bornean orangutan.

